# Development of a novel, non-invasive and whole brain biomarker of demyelination in a mouse model of multiple sclerosis

**DOI:** 10.1101/2024.03.18.585596

**Authors:** Benoit Beliard, Lauriane Delay, Isabelle Rivals, Youenn Travert-Jouanneau, Nathalie Ialy-Radio, Annabelle Réaux-Le Goazigo, Thomas Deffieux, Daniel P. Bradley, Mickael Tanter, Sophie Pezet

**Author notes:** Corresponding author: Sophie PEZET, Address: Institute of Physics for Medicine Paris, ESPCI Paris, 10 rue Oradour sur Glane, 75015, Paris, France.

## Abstract

Multiple Sclerosis (MS) is an autoimmune disease of the central nervous system (CNS), affecting 2.8 million people worldwide, that presents multiple features, one of which is demyelination. Although treatments exist to manage the condition, no cure has been found to stop the progression of neurodegeneration.

To develop new treatments and investigate the multiple systems impacted by MS, new imaging technologies are needed at the preclinical stage. Functional ultrasound imaging (fUS) has recently been demonstrated to robustly measure brain cerebral blood volume (CBV) dynamics as an indirect measure of neural activity. This study aimed at proposing a new biomarker of de- and/or re-myelination in a mouse model of MS induced by cuprizone. We demonstrate first that extended demyelination induces an increased hemodynamic response in the primary sensory cortex both spatially and temporally, which is consistent with fMRI data collected on MS patients. Second, using descriptors of the evoked hemodynamic response, we show that 3 of these descriptors allows the prediction of the level of myelin in the primary sensory cortex (p=5. 10^−5^) and the thalamus (p=6. 10^−6^). The development of such a non-invasive biomarker is crucial in the MS field as is provides an extremely useful tool for both disease follow-up and drug development.

**RESEARCH IN CONTEXT:** *Evidence before this study:* Multiple sclerosis (MS) is an autoimmune neurodegenerative disorder of the central nervous system. It is the most common cause of neurological disability in young adults, affecting approximately 2.8 million worldwide. While the field of studies in MS has been very active at identifying the neurobiological cellular and molecular mechanisms underlying MS progression, the number of new treatments has been very limited so far, due to several factors, such as the lack of robust and non-invasive biomarkers of myelin loss in longitudinal studies (measurements during the development of the disease). Unfortunately, quantification of myelin loss, (one of the key neurobiological markers of MS progression) is classically performed post-mortem on fixed tissues, preventing longitudinal studies. Longitudinal follow up of an indirect measure of myelin loss is possible, using magnetic resonance imaging. However, the small size of rodent brains poses a challenge for conventional imaging techniques, requiring the use of high field magnet to achieve the necessary sensitivity and resolution.

*Added value of this study:* In this study, using a sensitive neuroimaging technique, we developed a simple, non-invasive, predictive biomarker able to quantify the individual amount of myelin content consistently and accurately in brain structures in mice.

*Implications of all the available evidence:* The development of such a biomarker is extremely important for the MS field as it will accelerate the pre-clinical tests for drug efficacy. The benefits provided by our biomarker encompass: 1) Enhanced sensitivity in individually quantifying myelin content, providing a more comprehensive assessment across diverse brain regions 2) Speeding up the process of the discovery, by reducing the number of animals required per group and 3) It will also likely lead to new scientific outcomes, as many more structures will be studied (most teams and drug compagnies only study the demyelination at the level of the corpus callosum). Finally, from a clinical perspective, given the brain alterations observed in this animal model closely mirror those observed in early stages in patients with MS, we anticipate our biomarker, with minimal additional refinements, to be readily applicable in clinical settings.

## INTRODUCTION

Multiple sclerosis (MS) is an autoimmune neurodegenerative disorder of the central nervous system. It is the most common cause of neurological disability in young adults, affecting approximately 2.8 million worldwide [1]. Chronic inflammation, demyelination, gliosis and neuronal loss are hallmarks of the pathology, producing plaque formation and tissue destruction, in both white and grey matter [2]. Preclinical studies are crucial for understanding these mechanisms and for the development of new therapeutics. Magnetic Resonance Imaging (MRI) is largely used for MS diagnosis and follow up, but to date, it is not able to distinguish among MS disease courses. Novel imaging techniques, such as magnetization transfer ratio, diffusion inversion recovery, and diffusion tensor imaging are expected to fill this knowledge gap [3]. However, the spatial resolution of most neuroimaging modalities is not suited for preclinical studies using mouse models of MS. The only current solution to reducing this limitation is to use high-field scanners, but their high cost and lack of portability are prohibitive for many laboratories. As a result, the data is scarce and the lack of standardized MRI protocols makes it difficult to gather data from different labs, hampering a global understanding of MS mechanisms in animal models.

Ultrafast ultrasound imaging, on the other hand, does not have these limitations in preclinical studies. This relatively new imaging technology detects changes of cerebral blood volume with a great sensitivity [4,5], allowing functional studies in both anesthetized [6,7] and awake animals [8,9], the study of functional connectivity [10,11], of microvascular flux at a microscopic scale with a large field of view [12,13] and finally functional studies at a microscopic scale [14]. Its portability and lower cost make it a potentially widespread technology [15]. Due to the need of innovative and non-invasive biomarkers allowing the identification of MS mechanisms and the follow up of therapeutic strategies, this study aimed at investigating the alterations of cortical hemodynamic response induced by sensory stimulations in a widely used animal model of MS. Cuprizone (bis-cyclohexanone-oxalyldihydrazone, CPZ) is a copper chelator that has degenerative properties on oligodendrocytes, inducing a chronic (non-immune) demyelination of the brain [16–18]. Interestingly, its withdrawal allows to study natural remyelination and its fostering by pharmacological intervention. Due to the large sensory innervation of the whisker pad in rodents [19], we chose the barrel cortex as a model system in this study. Our study demonstrates long-lasting alterations of the hemodynamic response in cuprizone-induced demyelinated animals, and three descriptors of this evoked response can be synthetized into a biomarker of the level of demyelination.

## MATERIALS AND METHODS

### Animals

All experiments performed in this study complied with the French and European Community Council Directive of September 22 (2010/63/UE). They were also approved by the local Institutional Animal Care and Ethics Committees (#59, ‘Paris Centre et Sud’ project #2019-41). Accordingly, the number of animals in our study was kept to the minimum necessary. Using previously published data and preliminary data on the effect of cuprizone treatment, we performed a G-Power analysis (https://www.psychologie.hhu.de/arbeitsgruppen/allgemeine-psychologie-und-arbeitspsychologie/gpower) and found that N = 8 mice per group were required to detect statistical differences in our imaging experiments.

### Sex as a biological variable

Our study examined male mice only because male animals exhibited less variability in phenotype. It is unknown whether the findings are relevant for female mice. Therefore, this is one of the limitations of our study.

### Experimental design

Our experimental design followed the ARRIVE guidelines. It included 32 adult mice. One week after their arrival in the laboratory, the animals were randomly assigned to 4 different groups of equal size (8) and imaged at baseline (=D0). Following this imaging session, each group was treated as follows (Figure 1A):

- The control group received regular food for the whole duration of the study, i.e. 7 weeks. To enable the comparison to the treatment groups, the animals of this group were imaged at 3W, 5W and 7W after baseline.
- The group ‘early demyelination’ was intoxicated with 0,3% Cuprizone for 3 weeks and imaged at 3W prior to perfusion.
- The group ‘progressive demyelination’ was intoxicated with 0.3% Cuprizone for 5 weeks. The animals of this group were imaged at 3W and 5W to follow the changes of evoked hemodynamic response over the course of demyelination.
- In the ‘remyelination group”, the animals were treated with 0,3% Cuprizone for 5 weeks, after which they were given traditional food. These animals were imaged at 3W, 5W and 7W (Figure 1A).

**Figure 1:**
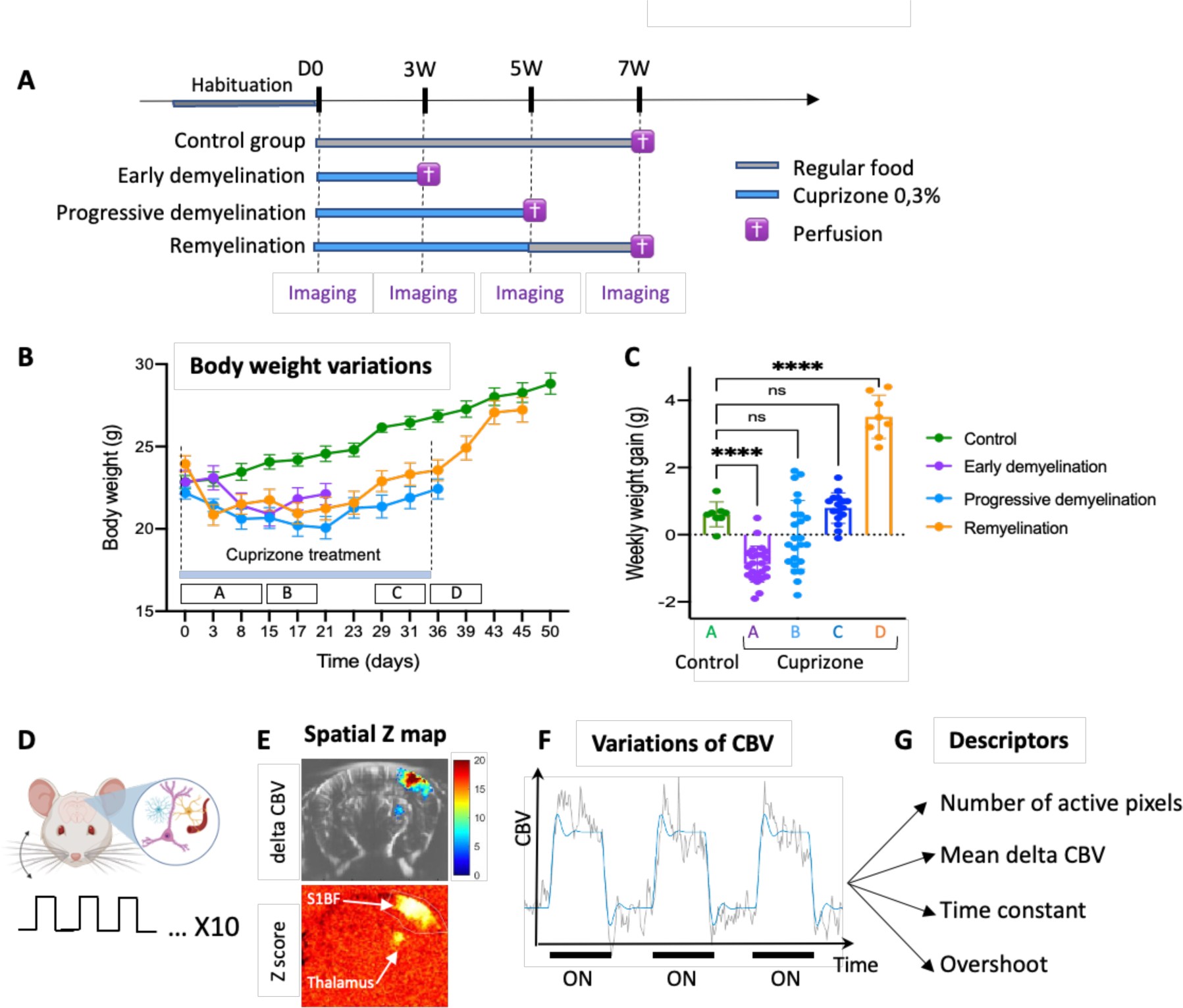
Study of functional hemodynamic response to whisker stimulation for the development of an innovative biomarker of demyelination. A: Experimental design. Four cohorts of 8 mice each were used in this study. These cohorts were: control (fed with regular food), ‘early demyelination’ (fed with cuprizone 0,3% for 3 weeks), ‘progressive demyelination’ (fed with cuprizone 0,3% for 5 weeks) and ‘remyelination’ (fed with cuprizone 0,3% for 5 weeks and then fed with regular food for 2 weeks). B-C: Variations of body weight (B) and weekly weight gain (C) in these different cohorts of animals showing the impairment of weight gain induced by cuprizone-induced demyelination and its recovery at discontinuation of the cuprizone treatment. Results are expressed as mean weight (g) or weekly weight gain (g) +/− SEM in N=8 animals per group. One-way ANOVA, followed by t-tests. **** p<0,0001. D: Study of the functional hyperaemia induced in the primary sensory cortex barrel field part (S1BF) by repetitive (10X) mechanical stimulation of the contralateral whiskers. Due to the increased neuronal activation and mechanism of neurovascular coupling, an increased cerebral blood volume is detected with ultrafast ultrasound imaging. First, Z score maps of activations were computed (E, bottom panel). E (top panel) shows an example of increased CBV in the S1BF and thalamus. After averaging the variations of CBV evoked extracted over the active pixels (defined as having Bonferroni corrected pvalue <0.05), in the S1BF (F), four descriptors were quantified: the number of active pixels, the steady state CBV variation ΔCBV, the time constant and the overshoot. The panel A was designed using Biorender.com.

As indicated in Figure 1, all animals were perfused intracardially at the end of the last imaging session in order to collect their brain and to perform immunofluorescent staining for Myelin Basic Protein (MBP). We did not exclude any animal.

The experimenter who performed the fUS imaging was blind to the animals’ treatment.

### Cuprizone administration

The experiments were performed on 32 male adult mice (C57BL/6 Rj, age 2-3 months, 20-30 g, from Janvier Labs, France). They were housed 8 per cage (also per group) in very large cages (11.818 in x 9.229 in x 16.228 in /300 mm x 234 mm x 412 mm, Floor space: 142 in^2^ | 916 cm^2^), in controlled conditions (22 ± 1°C, 60 ± 10% relative humidity, 12/12h light/dark cycle) and food and water a*d libitum*. Before the beginning of the experiments, the animals were acclimatized to the housing conditions for a minimum of one week.

Demyelination was induced in the three treatment groups by feeding the animals with a diet containing 0.3% cuprizone (bis-cyclohexanone oxaldihydrazone; Sigma-Aldrich Inc., St. Louis, MO) for 3 or 5 weeks, depending on the experimental group. Following (Zhan et al., 2020), in order to improve the reproducibility of cuprizone intoxication, 0.3g of fresh Cuprizone powder was mixed every day with 100g of ground food (SAFE, France). To follow the development of the model and to prevent dangerous weight loss, the mice were weighed twice a week. All cages were deprived of enrichment.

### Myelin immunofluorescent staining and its quantification

#### Perfusion

At the end of the imaging session, while the animals were still anesthetized, they received an intraperitoneal (IP) injection of 0,05 ml of Euthasol (Dechra Pharmaceuticals, concentration of the pentobarbital solution to 364,60 mg/mL). Then, a thoracotomy was performed, and an incision was made in the right atrium. Animals were perfused transcardially with 10mL of saline solution (0.9% NaCl), followed immediately by 40 mL of paraformaldehyde 4%. The brain was extracted and fixed overnight in 4% paraformaldehyde at 4°C before cryoprotection in a 30% sucrose solution for 2 days. They were frozen in an OCT (Optimal Cutting Temperature) matrix in cooled isopentane (−40°C) on dry ice and stored at −80°C until further process. The brains were cut in 12 μm slices on a cryostat (Leica CM 3050 S, Wetzlar, Germany), mounted on Superfrost slides (Thermofisher scientific, Waltham, Massachusetts, USA) and kept at −20°C in cryoprotectant until all brains from the whole cohort were collected (Pezet, 2002).

#### Myelin-Basic Protein (MBP) immunohistochemistry

Two slides (each one containing eight sections) of each animal were washed three times in 0.1M phosphate buffer with 0.9% NaCl (PBS). They were incubated overnight at room temperature with the primary antibody: mouse anti-MBP, clone 12 (Chemicon, Avantor-VWR, #MAB384, 1:150), diluted in 0.3% triton X-100. After the three washes, they were incubated for 2h with the secondary antibody (Alexa Fluor 488-conjugated donkey anti-mouse antibody, 1:1000; Invitrogen). Finally, the sections were cover slipped using Fluoromount (Sigma, Aldrich). The sections from all animals were stained simultaneously.

#### Microscopic observation and quantification

All stained sections were digitized using a nanozoomer (Zeiss, facility of the Vision Institute, Paris) using the same parameters of acquisition. The files obtained were then opened with the manufacturer’s software and two sections per animal, corresponding to sections at the coordinates: Bregma −1.34 mm and −1.70 mm, were exported in PNG format. The mean gray intensity of MBP staining was quantified on both the right and left sides of each section and averaged. Using Fiji software, after transformation in 256 grey level images, the mean gray intensities of the background (outside the brain, supplementary figure 2) and of the MBP staining in various brain regions were quantified (supplementary figure 2). After subtraction of the background level on each section, averaging over the right and left side, the values obtained from the two sections were finally averaged. The experimenter who performed the analysis was blind to the animals’ treatment.

### Imaging functional hyperaemia and quantification

#### Anesthesia

The mice were first induced with 2% Isoflurane and an intramuscular (IM) injection of medetomidine (Domitor, 0.08mg/kg). The anesthesia was then maintained with a continuous IM injection of medetomidine (0.08 mg/kg/h) using a syringe pump, with 0,5% isoflurane (0% O2, 100% air as carrying gas). The depth of anesthesia was monitored throughout the imaging session with the cardiac and respiratory frequencies (Labchart). The temperature of the mice was also monitored, and a heating pad was placed under the mouse to prevent hypothermia.

#### fUS Imaging

All the acquisitions were performed with the Iconeus One scanner (Iconeus, Paris, France) dedicated to small animals. The linear probe we used had 128 piezoelectric transducers with a 15 MHz central frequency. Ultrafast images were acquired at 500Hz using the coherent summation of 11 compounded tilted plane waves with angles separated by 2° and ranging from − 10° to 10° emitted at a 5500 Hz pulse repetition frequency.

The blood signal was then separated from the tissue signal thanks to the singular value decomposition clutter filter [20]. Each power Doppler frame is the average of 200 filtered ultrafast frames resulting in a 2.5Hz framerate.

Alterations such as scars or inflammations of the skin above the skull can lower the fUS signal due to reflections of the US waves at the interfaces. To remove such artifacts, at the beginning of each imaging session and under anesthesia, the skin was cut and opened below the US probe after local subcutaneous injection of Lidocaine. To prevent infections, the skull was covered by a sterile plastic film on which saline and 1 ml of ultrasound gel was dropped.

To improve the reproducibility of the imaging plane, the Icostudio software (Iconeus, Paris, France) was used to register its location on the Allen Brain Atlas after its alignment with the 3D vasculature of the mice measured by fUS [21]. The first step was to acquire a 3D scan (6mm span along the anterior-posterior axis, 0.2mm step) of the mouse brain to compute the coordinates of the 4-axis motor that will place the probe above the imaging plane set on the atlas.

#### Whisker stimulation

To further enhance the reproducibility of the measures, the whisker stimulation was standardized and controlled using an Arduino Uno (https://store.arduino.cc/; [22]). The system also allowed the synchronization of the stimulation (done by a Servomotor) and of the acquisition. The servomotor moved up and down a cotton bud placed perpendicularly to the whiskers at a frequency of 4Hz. After 20s of baseline, the stimulation pattern consisted of 20s of stimulation and 20s of rest, repeated 10 times (420s in total).

#### Signal analysis

After a detrending step to remove the slow signal drift occurring during the acquisition, a general linear model (GLM) analysis was performed to compute the activation maps displaying the significantly active area in color (Z scores with p-value < 0.0000006 = 0.05 divided by the number of pixels in the image using Bonferroni’s correction for multiple comparisons, Figure 1E, bottom panel). Quantitative information regarding the spatial aspect of the functional hyperemia was measured by counting the number of significant active pixels, thus quantifying the surface of activation. The average temporal signal was estimated on the active pixels and was then fitted through a continuous-time model using MATLAB’s function “idproc”. In the case of an underdamped response, a second order system with complex poles (P2U) characterized by its steady state gain ΔCBV, damping ζ and natural pulsation ω_n_ was used, and else a first (P1) or second order (P2) system with one or two real poles, as appropriate, characterized by ΔCBV and time constant τ (P1), or ΔCBV and time constants τ_1_ and τ_2_ (P2). For the three system types, a time constant (TC) was defined as (ζ ω_n_)^−1^ for P2U, as τ for P1 and as τ_1_ + τ_2_ for P2; the overshoot (OS) was defined as exp(– π ζ (1-ζ^2^)^−1/2^) for P2U, and as 0 for P1 and P2, see Supplementary Figure S1.

#### Statistical analysis

First, using GraphPad Prims 10, the normality of the data was tested using Shapiro-Wilk’s test and its homoscedasticity with Barlett’s test.

The following statistical analysis were performed using GraphPad Prism 10. The comparison of the mean grey density of MBP immunostaining between the various brain areas was performed using:

i) an ANOVA followed by Tukey’s multiple comparison test when the values followed a normal distribution.
ii) In the case of data that followed a normal distribution, but were heteroscedastic, we used the Brown Forsythe and Welch ANOVA test, followed by Dunnett’s T3 multiple comparison tests.

The statistical analysis of the four descriptors of the hemodynamic response measured by fUS imaging (i.e. number of active pixels, steady state CBV variation ΔCBV, time constant TC and overshoot OS) across time was performed using a repeated measures one-way ANOVA, followed by Tukey’s test for post-hoc multiple comparisons.

### Prediction using multiple linear regression

As the development of new pharmacological treatments requires a longitudinal measure of the level of myelin loss, this study aimed also at developing a quantitative surrogate for the level of myelin, through a longitudinal study of the altered hemodynamic response. To do so, we investigated the potential ability of the four descriptors of the hemodynamic response measured by fUS (number of active pixels, steady state CBV variation ΔCBV, time constant TC, overshoot OS) imaging to predict the level of myelin (quantified by MBP immunohistochemistry at the end of the experiment). To do so, we trained a multiple linear regression model with the four descriptors as inputs (note that 1/TC was used instead of TC). To assess the overall performance of the model, we conducted an F-test to compare the model with all its predictors to a constant model (analysis of variance test). Furthermore, we assessed the contribution of each predictor by testing whether the corresponding parameter significantly differs from zero using the classic t-test.

## Results

### Induction of myelin loss by Cuprizone treatment

While control animals had an approximately constant weekly weight gain of 0.61 g ± 0.13, the cuprizone treatment impacted the weight gain. But interestingly, this impact was not linear with time. From day 3 until day 15, animals lost weight on average (−0.88 g ± 0.19, Figure 1B, C). Between day 15 and day 21, some animals continued to lose weight, while others started gaining weight (averaged gain weight of 0.0 g ± 0.36 g over N=24 animals, figure 1C). Over the last week of cuprizone treatment (between 4 and 5 weeks of treatment (day 28-35)), the weekly gain weight returned to control values (+0.79 g ± 0.16, figure 1C). Finally, the reintroduction of standard food induced a highly and significantly increased weekly gain rate (+3.5 g ± 0.23). These results are in accordance with previous observations and suggest that the model developed as expected, affecting the natural ability of animals to gain weight during the course of demyelination [23].

### Measure of the evoked hemodynamic response at various stages of demyelination in the cuprizone model

Using functional ultrasound imaging, we quantified the putative alterations of the hemodynamic response in this model, in link with the demyelination evolution and / or the remyelination (5 weeks cuprizone + 2 weeks). To do so, we defined four descriptors of the brain hemodynamic response to a step mechanical stimulation of the whiskers: the number of active pixels, the steady state CBV variation, the time constant and finally the overshoot (Figures 1D, 1G, Supplementary figures 1, 3, 4, 5 and 6).

In the control cohort, we evaluated the stability and reproducibility of these measures over time in naïve untreated mice. Steady stateCBV variation, time constant and overshoot did not vary significantly with time (Figure 2A). The number of active pixels however was stable for the three first time point (DO, 3W, 5W), but significantly decreased at the latest time point (7 weeks), suggesting an effect of aging only at this latest stage. This effect is likely due to the thickening of the mouse skull during aging.

**Figure 2:**
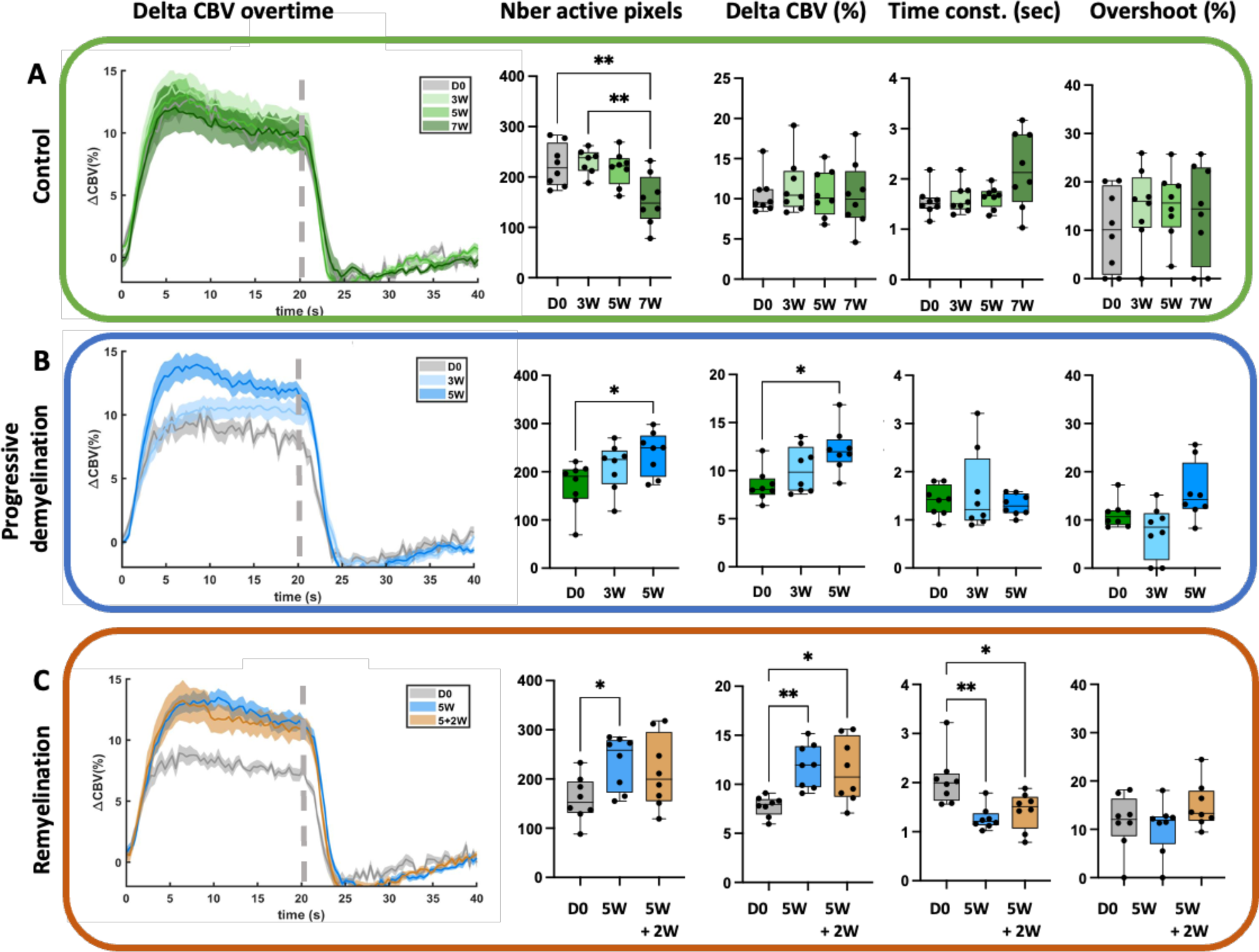
Longitudinal follow up of the cortical hemodynamic response during the course of demyelination induced by Cuprizone intoxication and its spontaneous (partial) remyelination. Mean relative changes of CBV evoked by whisker stimulation (curves in transparency define the SEM, n = 8 per group) in control animals (A), animals of the ‘progressive demyelination’ treated with cuprizone for 5 weeks (B) and of the ‘Remyelination’ groups, intoxicated for 5 weeks with cuprizone followed by 2 weeks of remyelination (C). In panels A, B and C, the 5 panels from left to right present for each cohort at different time points during the course of the disease: i) the mean time course of ΔCBV induced by whisker stimulation. The dotted lines represent the end of the stimulation (20 sec stimulation period). ii) The number of active pixels, iii) Steady state CBV variation ΔCBV (expressed in %, within the 7-20 sec stimulation interval), the time constant (sec) and finally the overshoot (%). In all panels, the data are presented as box plots, with an overly of individual values. As the data are individual animals followed over the course of the disease, we used a repeated measures one-way ANOVA, followed by Tukey’s test for post-hoc multiple comparisons for the statistical analysis. For esthetic reasons we are only showing the statistically significant groups: * p<0.05, ** p<0.001. N=8 animals per cohort.

To follow the putative alterations of functional hyperemia during demyelination, a second cohort (‘progressive demyelination’) was imaged before (D0), at early time point of demyelination (3W) and at maximal demyelination (5W). Our results show a progressive increase in the number of active pixels (p=0.03, Figure 2B), suggesting a more spread-out activation, and an increased CBV within these pixels (Figure 2B). These effects were statistically significant only at 5 weeks of cuprizone treatment. A decreased time constant and a slightly (non-statistically significant) decreased overshoot were observed, but only at 3 weeks of cuprizone treatment.

The remyelinating effects on the hemodynamic response were studied on a third cohort of animals (‘Remyelination’), imaged at D0, 5 weeks and after 2 weeks of normal food (5+2W). The effects of the 5 weeks of cuprizone treatment were like those described in the previous cohort, showing the reproducibility of our results across batches of animals treated with cuprizone. The effect induced by remyelination was modest and variable between animals, giving rise to more dispersed measures (Figure 2C). As will be discussed later, this variability in the hemodynamic responses is likely due to the inter-individual variability in the spontaneous remyelination between animals, as was raised by previous studies [24].

### Prediction of the level of demyelination using descriptors of the hemodynamic response in fUS imaging

The level of demyelination was assessed using MBP immunofluorescent staining in several areas of the brain. After 3 weeks of cuprizone treatment, which was described in many studies as an early time point in the demyelination process in this model [18,25,26], we observed that a strong and statistically significant demyelination occurred in both the primary sensory cortex barrel field part (S1BF) and the hippocampus (Figure 3 A, 3B, 3E and 3G). Surprisingly, an increased MBP immunostaining was observed in the corpus callosum at this time point (Figure 3F).

**Figure 3:**
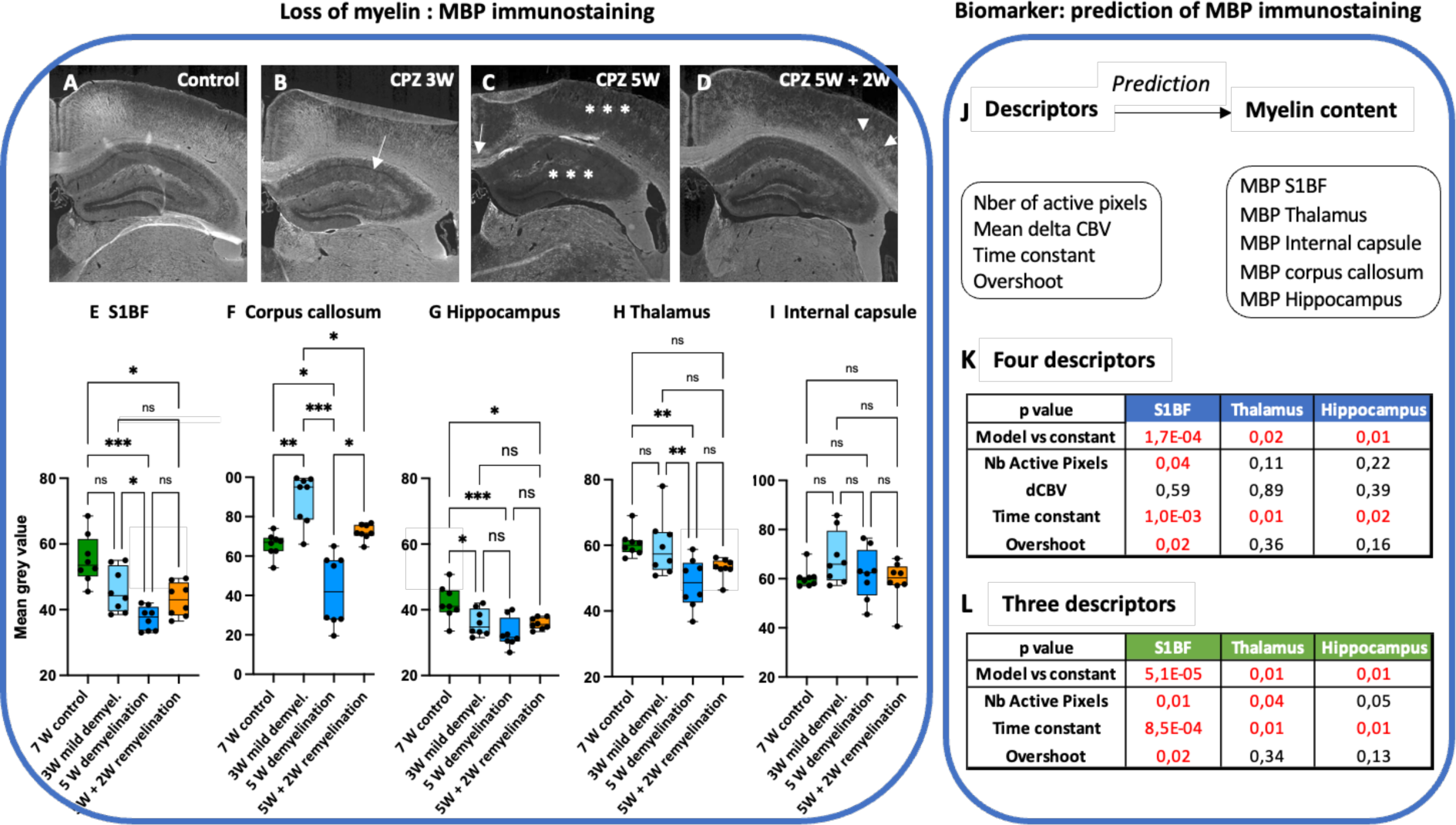
Use of four descriptors extracted from the evoked hemodynamic response to predict the level of Myelin of individual animals. A-D show representative examples of MBP immunostaining in the S1BF cortex, corpus callosum and hippocampus in control animals (A) and animals intoxicated with Cuprizone (CPZ) for 3W (B), 5W (C) or let to remyelinate 2 weeks after the 5 weeks of cuprizone intoxication (D). Arrow in B shows the beginning of demyelination in the hippocampus. In C, the arrow shows a strong demyelination in the corpus callosum (arrow) and the cortex (stars). Arrows in D point at the patches of remyelination in the S1BF. E-I show the quantified MBP immunostaining and the loss of myelin induced by CPZ treatment in the S1BF (E), medial corpus callosum (F), hippocampus (G), and thalamus (H) and the complete or partial remyelination depending on the brain area. At the time point studied, there was no demyelination in the internal capsule (I). Results are presented as boxplots, with an overlay of individual values. Statistical analysis used: ANOVA followed by Tukey’s multiple comparison test when the values followed a normal distribution and were homoscedastic. In the case of normality and heteroscedastic data (S1BF, hippocampus and internal capsule), the Brown Forsythe and Welch ANOVA test was used, followed by Dunnett’s T3 multiple comparison tests. * p<0.05, ** p<0.01, *** p<0.001. NS: Non-significant. N=8 per group. Scale bar: 1mm. J-L: Use of descriptors extracted from the evoked hemodynamic response to predict the level of Myelin of individual animals in various brain areas. J: Using the four descriptors of the hemodynamic response from figure 2, we used linear models in order to predict the level of MBP in the brain areas analyzed above. K and L report the p-values (corresponding to the null hypothesis that the estimate is equal to zero) of the models and descriptors for the model trained, using either 4 (K) or 3 (L) descriptors. As the model in K shows that the ΔCBV is not statistically significant for the model, the model in K was run without it. Results of both models show consistently that measurements of the number of active pixels, the time constant and the overshoot are key element for the prediction of the myelin content in the S1BF. The time constant is also a very valuable measurement for the prediction of the MBP in the thalamus and hippocampus, as well as the number of active pixels for the prediction of the MBP in the thalamus.

As expected, 5 weeks of cuprizone treatment induced a strong demyelination in many brain areas, such as the S1BF (Figures 3C, 3E), the medial corpus callosum (Figures 3C, 3F), the hippocampus (Figures 3C, 3G) and the thalamus (Figure 3I). Switching to normal food for two weeks induced remyelination in the four aforementioned areas (Figures 3D-H).

Interestingly, the internal capsule was subjected neither to demyelination, nor to remyelination (Figure 3J).

Finally, we assessed the predictive capacity of the four descriptors of the hemodynamic response to determine the level of myelin in five brain regions using linear regression models. For each brain region where MBP was quantified, a linear model was trained using the four descriptors (Figure 3J). As reported in Figure 3K (and in supplementary table 1), the models accurately predicted the MBP levels only in the primary sensory cortex (p-value = 1.6 10^−4^), thalamus (p-value = 1.8 10^−2^) and hippocampus (p-value = 1.1 10^−2^). To further investigate the most significant predictors, we examined the estimate and corresponding p-value of the parameter of each predictor in the model predicting the MBP in the median corpus callosum (Figure 3K). Interestingly, the steady state CBV variation ΔCBV was never a significant descriptor. The time constant was however a significant descriptor in the prediction of the myelin loss in these brain areas. The overshoot and the number of active pixels were significant descriptors of the extent of demyelination in the S1BF.

As the ΔCBV was not an important descriptor for the prediction of MBP content, we next generated a second series of linear models without ΔCBV (three descriptors, Figure 3L). In agreement with the previous analysis, the models obtained (figure 3L) predicted also the MBP levels in the same areas, but with stronger statistical significance: primary sensory cortex (p-value = 5.0 10^−5^), thalamus (p-value = 6.8 10^−3^) and hippocampus (p-value = 5 10^−3^). In these models, the three descriptors were necessary for the prediction of the MBP in the S1BF, the number of active pixel and the time constant for the prediction of the MBP in the thalamus, whereas the time constant was the only significant descriptor for the prediction of the MBP loss in the hippocampus.

As reported in Supplementary figure 7, none of the models succeeded in predicting the myelin loss in the corpus callosum and internal capsule.

## DISCUSSION

Due to the large need of non-invasive methods to quantify myelin content in preclinical models of MS and the high sensitivity of fUS imaging towards changes of cerebral blood volume, this study investigated the alterations of whisker stimulus-evoked hyperemia in the cuprizone model in mice. Using four descriptors of this altered hemodynamic response, we could build novel biomarkers of the demyelination, predictive of the content of myelin in the primary sensory cortex, thalamus, and the hippocampus.

### Clinical relevance of the increased evoked hemodynamic response observed in the cuprizone model

To study the changes of evoked response associated with the different phases of the cuprizone model, we made the choice to perform a longitudinal study; each animal being its own control. Our results show that in the two cohorts where animals were treated for 5 weeks with cuprizone, the whisker stimulus-evoked hyperemia was consistently increased. The activated zone was larger (a larger number of active pixels) and the activation was enhanced in this zone (an increased CBV over time).

These results are highly relevant to the clinical condition [27], as in MS patients with clinically isolated syndrome (CIS [28,29], with non-disabling [30] or mildly disabling relapsing-remitting (RR) MS [31], a motor task induces an increased activation of the primary sensorimotor cortex. These changes that were confirmed in multiple centers (see for review [32]) have been considered to play a compensatory role in RRMS, since it was related to maintaining a good task performance despite the presence of widespread structural damage, and are believed to be due to both altered cerebral blood flow and oxygen consumption [33,34].

This interesting parallel between our results and the clinical condition suggests first, that even though the cuprizone model does not model all hallmarks of the clinical condition, it modeled well its altered hemodynamic responsiveness. Several authors have hypothesized that these surprising, enhanced responses might be an adaptive plasticity to compensate for the growing disability of the diseased brain. The cellular and molecular mechanisms behind this plasticity are still largely unknown. There is, however, a large literature describing an alteration of the vascular system and of the neurovascular unit in MS [35], opening the debate on whether vascular events may be the primary cause of neurological diseases or rather a mere participant recruited from a primary neuronal origin. Today, our results suggest that demyelination, induced by oligodendrocyte disruption, is able by itself to promote such alteration of the neurovascular coupling.

One of the limitations of our study, is the lack of investigation of the extent of this neurovascular alteration at later time points (10-12 weeks of cuprizone treatment), which is known to mimic some aspects of the late stage of clinical MS, characterized by axonal damage. Such functional study would reveal if, as observed in MS patients, the functional response is decreased. This will be the focus of future work.

### Cuprizone induced inhomogeneous brain demyelination

In order to develop this new biomarker, we quantified in our cohorts of mice the content of myelin, through the mean gray density of the Myelin Basic Protein (MBP) immunoreactivity. Most teams only follow up the level of demyelination in the medial corpus callosum [18,25,36]. Interestingly, as reported in some studies, there is a large variability in the extent of myelin loss between the various cerebral structures [37]. The internal capsule was not demyelinated after 5 weeks of cuprizone treatment, which is consistent with previous observations [37]. The S1BF, hippocampus and medial corpus callosum were the first structures to lose myelin (as soon as 3 weeks of treatment), followed by the thalamus (5 weeks only), which is consistent with previous studies [36–41]. Surprisingly, we observed a consistent increased MBP immunoreactivity in the corpus callosum at 3 weeks, which is contradictory with the well described demyelination of the corpus callosum that starts at 3 weeks according to several authors [36] and peaks at 5 weeks. Our interpretation of this surprising result is that, as proposed by Plemel et al., at early time in the model development, as myelin sheath is being attacked by pro-inflammatory response, the damaged myelin is more accessible to antibodies, leading to an increased immunoreactivity [42]. The question remaining is ‘why is it the case in the corpus callosum and nowhere else?’. The answer might be that it is among the most abundant bundles of myelinated fibers. This might explain why, despite the lack of statistical changes in the MBP level in the internal capsule (another structure particularly rich in myelinated fibers), our quantification shows a large variability with an increased average content compared to control mice (Figure 2J).

### New biomarkers of the level of myelin in the S1BF, thalamus and hippocampus

As sensory afferents from the whiskers go through the SpV nucleus to the thalamic nuclei VPM and Po [43] and finally the information conveyed by the thalamo-cortical neurons run through the internal capsule [44], it was legitimate to hypothesize that the alteration of evoked cortical response in the S1BF to whisker stimulation might be associated with myelin loss in some parts this sensory pathway, i.e. the thalamus, internal capsule and / or the S1BF. Indeed, our results confirm this hypothesis: the abnormal hemodynamic response in the S1BF is predictive of the loss of myelin in both the S1BF and the thalamus. This was not the case in the internal capsule, probably due to the lack of significant demyelination at these time points.

Interestingly, the key elements of this biomarker did not include the ΔCBV (the amplitude of the response), but the increased spatial response (increased number of active pixels) and the delay of this response (decreased time constant) in Cuprizone-treated animals. In the S1BF, we also observed a significant involvement of the overshoot in the prediction of the myelin loss. The overshoot reflects an exacerbated vascular response compared to the steady state CBV, in sustained (20-40 sec) stimulations [45]. It is a large vascular response that exceeds the physiological demand, a mechanism largely observed in the mechanism of neurovascular coupling [46], and yet whose physiological role is a debate [47,48]. Modeled by several teams [45,49], the overshoot is observable in the balloon model when the viscoelastic time constant is non-zero [49]. Only a few studies have determined experimentally the physiological factors influencing the overshoot. It is accepted that its initial trigger is the neuronal electrophysiological activity [50] and the overshoot of oxygen during neurovascular coupling depends upon the pO_2_ at the baseline (i.e. before the stimulation); a mechanism believed to be due to a sustained drop of oxygenation in cerebral tissues located remotely from the vascular feeding sources [51]. While there is accumulating evidence that there is some hypoxia in MS patients [52] and in the EAE model of MS [53], according to Hashem and coll., there is none in the cuprizone model [54]. Further work on the neurobiology of the overshoot in this model and in the clinical condition is required to finally clarify by which mechanism the overshoot is such a strong indicator of demyelination in the cuprizone model.

## Author contributions

SP, BB, MT and DPB designed the experimental paradigm.

NIR and BB performed the ultrasound experiments and BB processed the ultrasound data. TD and MT supervised the signal processing of the ultrasound data. YT performed the immunohistochemical experiments, while LD and ARL performed their acquisition and quantifications. BB and SP wrote the manuscript.

IR and BB assessed and verified the data, performed the statistical analysis, and wrote the corresponding section in the manuscript.

BB, SP, MT and IR were involved in the interpretation of the data and wrote some parts of the manuscript.

All authors read and corrected the final version of the manuscript.

## Acknowledgments

This work was supported by Biogen (B. Beliard’s PhD fellowship), the ANRT (B. Beliard’s PhD fellowship), the Institut National de la Santé et de la Recherche Médicale, Biogen and the European Research Council (ERC) Advanced Grant FUSIMAGINE and the ESPCI.

Authors would like to thank Mrs L. Antoinette for animal husbandry and Miss Celia Amabile for her early involvement in the project. Finally, this work was supported by the Inserm ART (Technology Research Accelerator) in “Biomedical Ultrasound”.

Several panels of figures were made using Biorender (Biorender.com).

## Data availability statement

Source data are available on the repository website Dryad using the following link: 10.5281/zenodo.10518303.

## Code availability statement

Custom codes used for the analysis of fUS data used in this study are protected by INSERM. The rest of this work did not involve any other particular code.

