## Supplementary figures for "Development of a novel, non-invasive and whole brain biomarker of demyelination in a mouse model of multiple sclerosis"

**Supplementary materials**

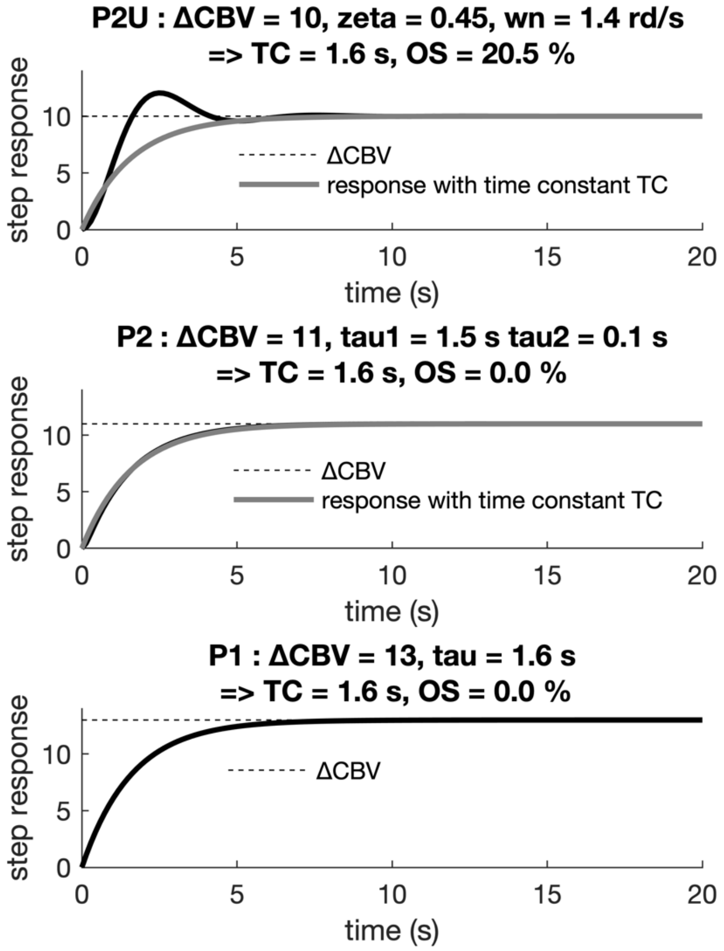


**Supplementary figure 1: Schematics illustrating how the steady state CBV variation, the time constant and the overshoot were determined from evoked hemodynamic response induced by whisker stimulation depending on the identified dynamic model.** The temporal signal then fitted through a continuous-time model using MATLAB's function "idproc". In the case of an underdamped response, a second order system with complex poles (P2U, as in A) characterized by its steady state gain ΔCBV, damping $\zeta$ and natural pulsation $\omega_n$ was used, and else a first (P1, as in C) or second order (P2, as in C) system with one or two real poles, as appropriate, characterized by ΔCBV and time constant $\tau$ (P1), or ΔCBV and time constants $\tau_1$ and $\tau_2$ (P2). For the three system types, a time constant (TC) was defined as $(\zeta\ \omega_n)^{-1}$ for P2U, as $\tau$ for P1 and as $\tau_1 + \tau_2$ for P2; the overshoot (OS) was defined as $\exp(-\pi\ \zeta\ (1-\zeta^2)^{-1/2})$ for P2U, and as 0 for P1 and P2.

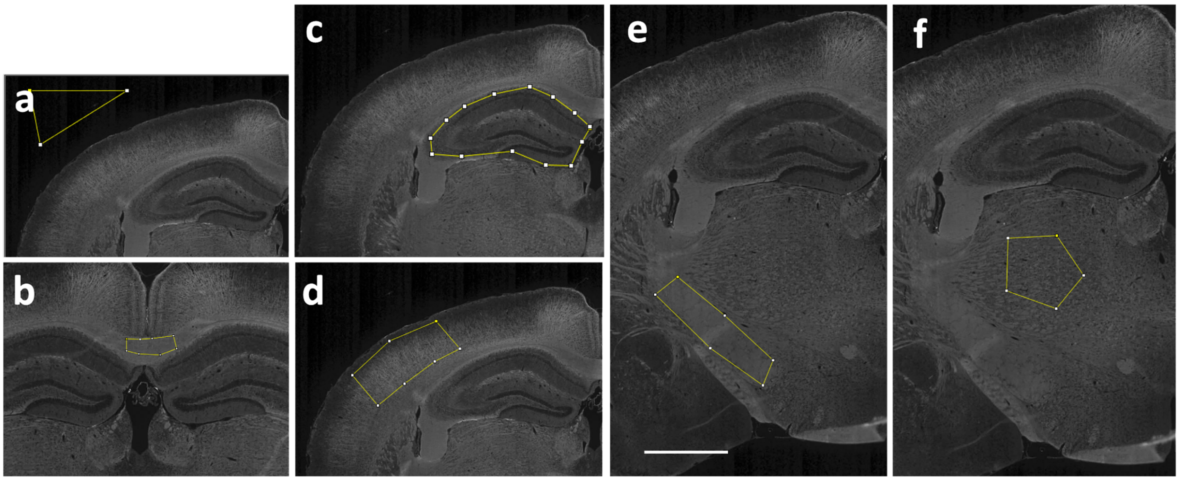


**Supplementary figure 2: Contour of brain areas where MBP immunostaining was quantified.** Each picture illustrates the area taken to quantify the mean grey value on each section in the background (a), medial corpus callosum (b), hippocampus (c), S1BF (d), internal capsule (e) and thalamus (f).

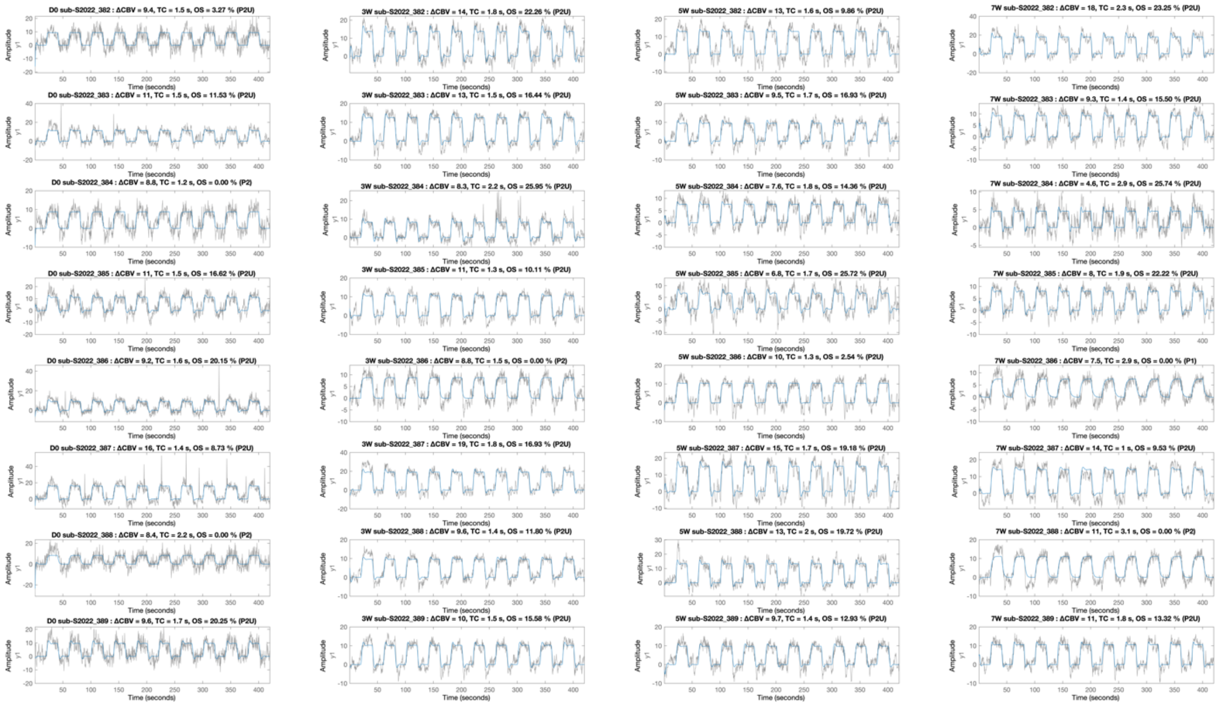


**Supplementary figure 3: Reporting of individual traces of CBV over time for all the animals of the cohort of control animals, at the four time points studied (D0, 3W, 5W and 7W).** Each column is a time point. Results from each animal are presented in lines. Each graphic shows the CBV variations over time (10 whiskers stimulations). The various descriptors measured from the graphic are presented on top of it, as well as the type of model (P2U, P2 or P1).

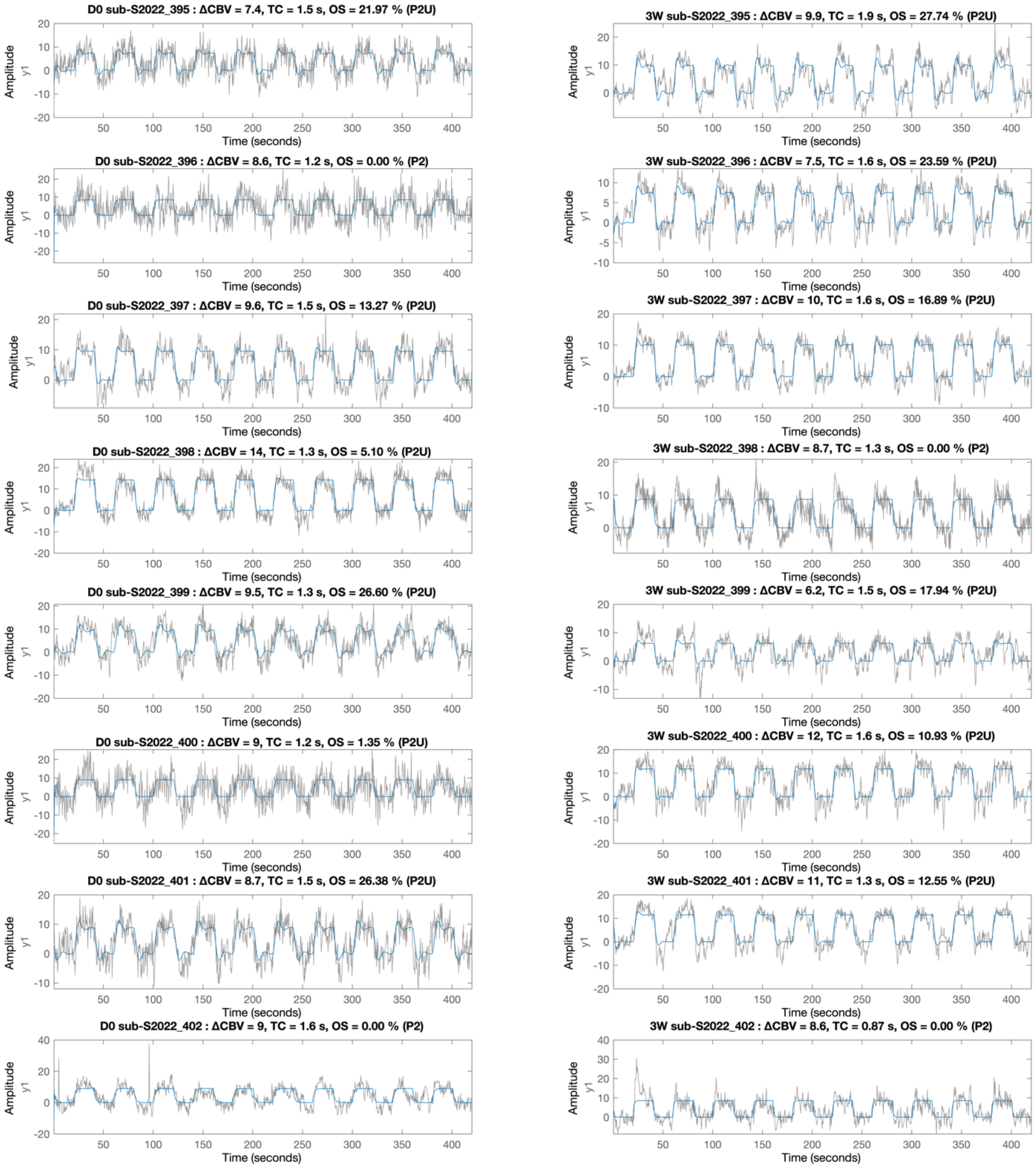


**Supplementary figure 4: Reporting of individual traces of CBV over time for all the animals of the cohort 'early demyelination', at the two time points studied (D0 and 3W).** Each column is a time point. Results from each animal are presented in lines. Each graphic shows the CBV variations over time (10 whiskers stimulations). The various descriptors measured from the graphic are presented on top of it, as well as the type of model (P2U, P2 or P1).

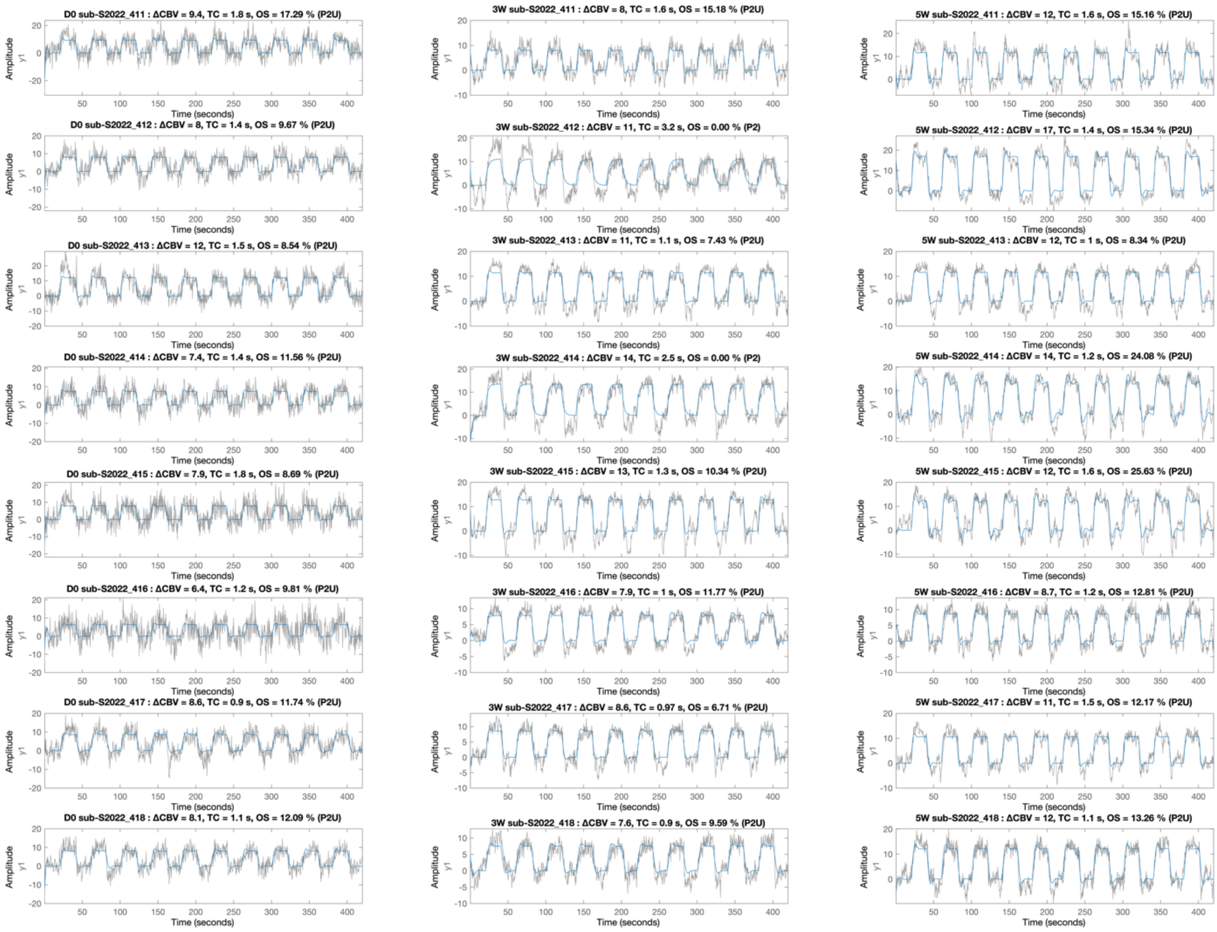


**Supplementary figure 5: Reporting of individual traces of CBV over time for all the animals of the cohort 'progressive demyelination' at the three time points studied (D0, 3W and 5W).** Each column is a time point. Results from each animal are presented in lines. Each graphic shows the CBV variations over time (10 whiskers stimulations). The various descriptors measured from the graphic are presented on top of it, as well as the type of model (P2U, P2 or P1).

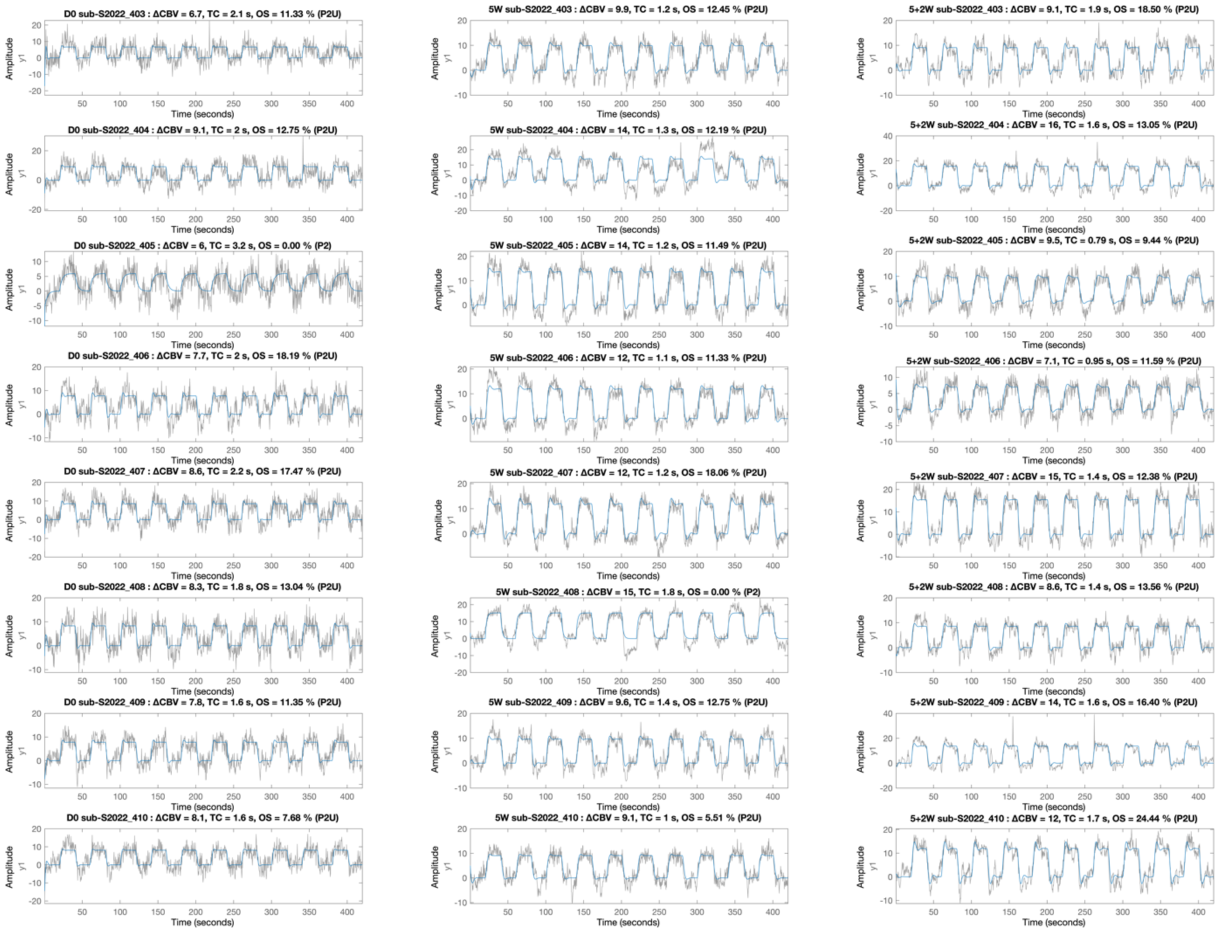


**Supplementary figure 6: Reporting of individual traces of CBV over time for all the animals of the cohort 'remyelination' (5W + 2W), at the three time points studied (D0, 3W, 5W and 7W).** Each column is a time point. Results from each animal are presented in lines. Each graphic shows the CBV variations over time (10 whiskers stimulations). The various descriptors measured from the graphic are presented on top of it, as well as the type of model (P2U, P2 or P1).

**A** Four descriptors

| p value | S1BF | Internal Capsule | Thalamus | Corpus Callosum | Hippocampus |
|---|---|---|---|---|---|
| Model vs constant | 1,67E-04 | 0,71 | 0,02 | 0,26 | 0,01 |
| Normality | 0,05 | 0,07 | 0,18 | 0,53 | 0,04 |
| Nb Active Pixels | 0,04 | 0,65 | 0,11 | 0,08 | 0,22 |
| dCBV | 0,59 | 0,76 | 0,89 | 0,88 | 0,39 |
| Time constant | 1,01E-03 | 0,26 | 0,01 | 0,54 | 0,02 |
| Overshoot | 0,02 | 0,66 | 0,36 | 0,57 | 0,16 |

**B** Three descriptors

| p value | S1BF | Internal Capsule | Thalamus | Corpus Callosum | Hippocampus |
|---|---|---|---|---|---|
| Model vs constant | 5,07E-05 | 0,56 | 0,01 | 0,15 | 0,01 |
| Normality | 0,07 | 0,09 | 0,21 | 0,56 | 0,07 |
| Nb Active Pixels | 0,01 | 0,72 | 0,04 | 0,03 | 0,05 |
| Time constant | 8,52E-04 | 0,25 | 0,01 | 0,53 | 0,01 |
| Overshoot | 0,02 | 0,67 | 0,34 | 0,55 | 0,13 |

**Supplementary figure 7:** Use of four (A) or three (B) descriptors extracted from the evoked hemodynamic response predict the level of Myelin of individual animals in the S1BF, thalamus and hippocampus. This figure is the extended version of the figure 3k, 3L.
